# Optogenetic Modulation of Cortical Neurons Using Organic Light Emitting Diodes (OLEDs)

**DOI:** 10.1101/669986

**Authors:** Arati Sridharan, Ankur Shah, Swathy Sampath Kumar, James Kyeh, Joseph Smith, Jennifer Blain-Christen, Jit Muthuswamy

## Abstract

**Objective:** There is a need for low power, scalable photoelectronic devices and systems for emerging optogenetic needs in neuromodulation. Conventional light emitting diodes (LEDs) are constrained by power and lead-counts necessary for scalability. Organic LEDs (OLEDs) offer an exciting approach to decrease power and lead-counts while achieving high channel counts on thin, flexible substrates that conform to brain surfaces or peripheral neuronal fibers. In this study, we investigate the potential for using OLEDs to modulate neuronal networks cultured *in vitro* on a transparent microelectrode array (MEA) and subsequently validate neurostimulation *in vivo* in a transgenic mouse model.

**Approach:** Cultured mouse cortical neurons were transfected with light-sensitive opsins such as blue-light sensitive channel-rhodopsin (ChR2) and green-light sensitive chimeric channel-rhodopsin (C1V1tt) and stimulated using blue and green OLEDs (with 455 and 520 nm peak emission spectra respectively) at a power of 1 mW/mm^2^ under pulsed conditions.

**Main results:** We demonstrate neuromodulation and optostimulus-locked, single unit-neuronal activity in neurons expressing stimulating and inhibiting opsins (n=4 MEAs, each with 16 recordable channels). We also validated the optostimulus-locked response in a channel-rhodopsin expressing transgenic mouse model, where at least three isolatable single neuronal cortical units respond to OLED stimulation.

**Significance:** The above results indicate the feasibility of generating sufficient luminance from OLEDs to perform neuromodulation both in vitro and in vivo. This opens up the possibility of developing thin, flexible OLED films with multiple stimulation sites that can conform to the shape of the neuronal targets in the brain or the peripheral nervous system. However, stability of these OLEDs under chronic conditions still needs to be carefully assessed with appropriate packaging approaches.

## INTRODUCTION

Organic light-emitting-diodes (OLEDs) are an emerging technology that can potentially be used for optogenetic applications. Optogenetics is a powerful tool for understanding the microscale spatial and temporal behavior of neural circuits that is applicable for brain mapping and for understanding mechanisms of neuronal dysfunction in debilitating diseases like epilepsy, Alzheimer’s, Parkinson’s, Huntington’s, and others [1]. Other applications include potential therapeutic interventions for localized modulation of seizure activity [2], stimulation of virally transfected retinal cells in retinitis pigmentosa using spatio-temporally targeted optogenetic stimulation [3]. The primary principle of optogenetics is to transfect a target neuron to genetically express a light-sensitive ion channel (such as channel-rhodopsins, halo-rhodopsins, bacteriorhodopsins etc.) that can be selectively stimulated or silenced with an external light source for localized neuromodulation [4][5]. Achieving spatio-temporal selectivity is partly dependent on the light source and its ability to selectively stimulate localized regions of the neural circuit. Typical light sources used are high-powered, high intensity laser LEDs, halogen lamps that deliver light via fiber optics [6][7][8]. Next generation light sources such as inorganic LEDs (GaN) have made considerable strides in improving power consumption, although high power consumption, and potential heat generation remain key issues [9].

In this study, we explore the potential for using OLEDs to electrically stimulate neuronal networks cultured in vitro on a transparent microelectrode array and in vivo in rodent experiments. OLEDs are low-power devices consisting of an electroluminescent stack of thin organic films sandwiched between two conductors acting as anode and cathode. Under forward bias conditions of the diode, OLED films luminesce. There are several inherent advantages of OLEDs compared to conventional LEDs that are appealing to an implantable neurostimulation approach. The flexibility of OLED thin films allows them to be readily placed adjacent to neuronal fibers in the peripheral nervous system or neurons of interest in the central nervous system with a potential to minimize foreign body response under chronic conditions. Their lateral dimensions can range from millimeters to tens of microns making them suitable for a range of neuromodulation needs from single cells to whole regions of the brain. The planar nature of the technology also lends itself readily to the patterning of multiple OLEDs with different wavelengths spatially placed according to the needs of the targeted neuronal network. In addition, arrays of OLEDs operating under active matrix conditions such as those operated in this study can significantly reduce power consumption, and lead-counts for interconnects. For instance, mxn arrays of OLEDs under active matrix conditions with thin film transistors can be operated with m+n leads as opposed to mxn leads. Other advantages include increase in brightness by tandem stacking without changes in current density. The diffuse light intensity is also more evenly distributed allowing for larger areas of stimulation. The OLED film uses an organic layer as the photosensitive hole-transport layer, where small molecule or polymer based emissive layers are typically used [10][11][12]. Unique materials such as nucleic acids have also been used to enhance performance [13]. OLED based systems are also used in biotechnology applications such as biosensors and as sources in protein microarrays [14][15].

One of the early studies on neurostimulation, used an 852 x 600 pixel OLED micro-display screen that was focused on an isolated retina using one aspheric lens to achieve pixel sizes of 4 μm × 4 μm and a mean light intensity of 8.6 nW/mm^2^. The study demonstrated neuronal responses in the form of spikes from ganglion cells [16]. Recently, OLEDs with a thin (1 μm thick) encapsulation layer for protection and at low power were used to modulate phototactic behavior of *Chlamydomonas Reinhardtii* and subsequently millimeter scale, higher power OLEDs (0.25-0.4 mW/mm^2^) triggered ChR2 induced muscle contractions in drosophila larvae [17][18]. A very recent study also demonstrated successful excitation of dose-dependent calcium transients in cultured cortical neurons expressing ChR2 using blue OLEDs [19]. However, the ability to stimulate action potential events opto-genetically (with opsins engineered for different wavelengths, sensitivities and reaction times) in neurons will make the OLEDs more widely applicable for neuromodulation studies.

The challenges in using OLEDs for neuromodulation involving neurons expressing opsins include the low luminance achievable with current OLEDs and the susceptibility of OLEDs to degradation in the presence of water and oxygen. These two significant barriers are not insurmountable with rapid advances in both OLED efficiencies and sensitivity of opsins. In the current study we operate the OLEDs in pulsed mode to achieve a maximum of 1 mW/mm^2^ output. We demonstrate modulation of action potential events in response to optical stimulation (by OLEDs) in rodent/mammalian neurons expressing opsins for excitation and inhibition in vitro and for the first time in vivo in transgenic mice. We use heat sinks and moisture barriers on the OLEDs to minimize ingress of moisture and oxygen. The fabrication of these OLEDs is detailed elsewhere [20]–[22] and the focus of this study is to assess the neuromodulatory effect of OLEDs on neural activity of cultured primary neuronal networks transfected with light-sensitive opsins such as blue-light sensitive channel-rhodopsin (ChR2) and engineered, red-shifted channel-rhopdopsin chimera (C1V1tt) that is sensitive to green-light. We demonstrate optostimulus-locked, single unit-neuronal activity for excitatory and inhibitory opsins on cultured cortical neurons on MEAs. We also validate the optostimulus-locked response in a channel-rhodopsin transgenic mouse model. The stability of such OLED based neurostimulation under long-term implantation conditions will be the focus of a future study.

## 2. MATERIALS & METHODS

### 2.1 OLED Fabrication and characterization

OLED fabrication is detailed elsewhere [20], [21]. Briefly, a thin-film, multi-layer device was constructed with a hole injection layer (HIL), hole transport layer (HTL), either a fluorescent blue [21] or phosphorescent green [22] layer, a hole-blocking layer (HBL), a doped electron transport layer, and electron injection layer. The above layers were sandwiched between a reflective aluminum cathode at the back end and a transparent indium tin oxide (ITO) under the HIL as shown in Fig. 1C. Blue and green OLED sources (4 mm^2^ electrode dimensions) were used in this study. OLEDs were fabricated at temperatures less than 200°C on 125 μm thick Dupont Teijin films Teonex poly-ethylene naphthalate (PEN) films bonded to rigid carrier. Additional moisture barrier layers were added to protect both the ITO anode and the aluminum cathode. After the thin film deposition processes are completed, the PEN substrate and the nanoscale thin OLED films deposited on top were peeled off from the rigid carrier.

**Fig.1.**
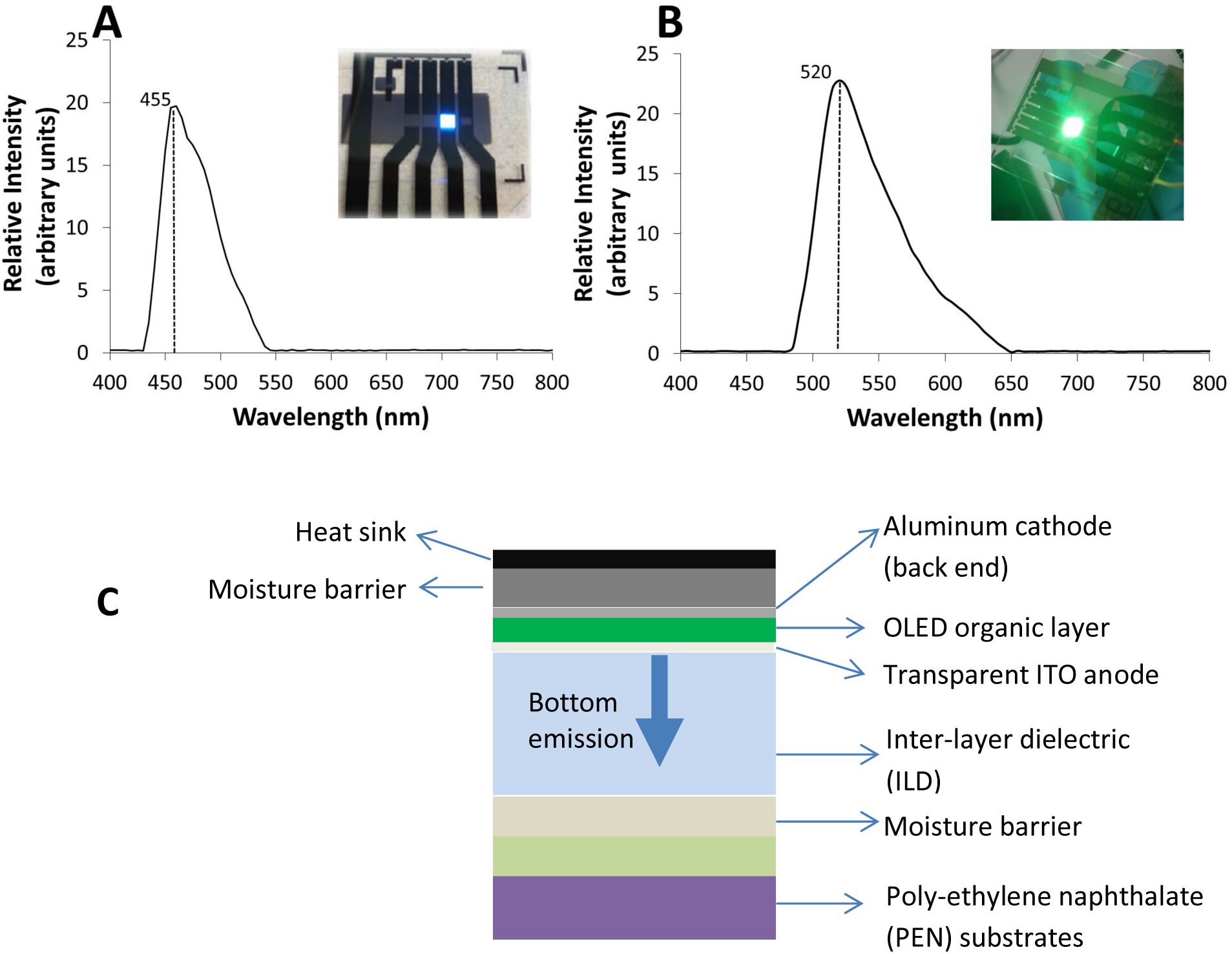
OLED emission spectra for (A) Blue OLED with wavelength centered at 455 nm and (B) green OLED with wavelength centered at 520 nm. The illuminated area in both cases is 4 mm^2^ (2 mm × 2 mm). (C) Schematic (not to scale) of the cross-section of the layers in the OLED coupon with the stack of OLED layers (shown in green) sandwiched between a reflective aluminum cathode on top (back end) and a transparent Indium tin oxide (ITO) anode. With subsequent layers of ILD (inter-layer dielectric) and a moisture barrier, the entire assembly is fabricated on 125 μm thick flexible, transparent PEN substrates.

The emission spectra of these OLEDs are shown in Fig.1, where blue OLEDs had a peak emission at 455 nm, while green OLEDs had a peak emission at 520 nm and the relative intensity for both colors were similar. For both in vitro and in vivo experiments, blue and green OLEDs were pulse-driven at 14V, 10 Hz at 10% duty cycle to achieve at least 1 mW/mm^2^ intensity. Fig. 2 shows the optical power density as a function of voltage bias for blue (Fig. 2a) and green (Fig. 2b) OLED coupons.

**Fig.2.**
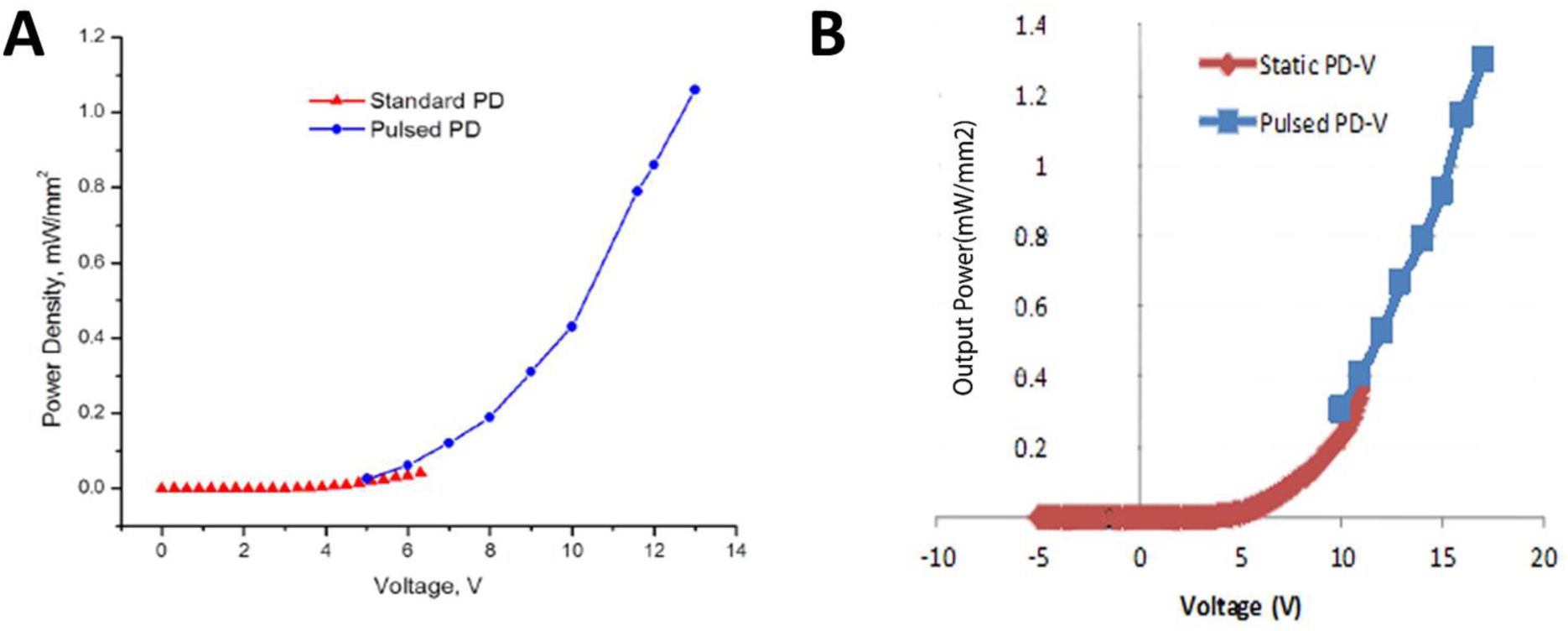
Optical power density as a function of voltage bias for (A) blue OLEDs and (B) green OLEDs. The red curve represents a DC voltage bias and the blue curves represent a 10 Hz pulse train (10% duty cycle) at the given voltage. The pulse-driven OLED for both blue and green varieties gave higher optical power density.

### 2.2 In vitro experiment setup

A typical experimental setup utilized a DC power supply that is modulated by a trigger signal from a function generator and a relay (Hamlin HE721C0510) to generate a 14V square pulse waveform with a 10% duty cycle (Fig.3). Pulses with a 10 msec pulse-width were delivered to OLEDs, which were then used to stimulate either transfected neurons in culture on MEAs or into the cortex of a channel-rhodopsin transgenic mouse. In vitro experiments utilized cortical tissue (E18) from Sprague-Dawley rats (Brainbits, Springfield, IL) seeded on custom-fabricated, transparent microelectrode arrays (MEAs) with 100 µm diameter electrodes as previously described in [23] (Fig.4). Briefly, MEAs were autoclaved and surface treated with 1 mg/ml polyethyleneimine (PEI) for 5 hrs, washed with distilled water once and dried for another hour. Neurons were dissociated and seeded on 4 different MEAs at ~3000 cells/mm^2^ over the electrode area and cultured in NbActiv1™ medium for at least 7-14 days prior to transfection with Fubi-ChR2-GFP (Addgene plasmid # 22051) or 21 days prior to transfection with C1V1tt (Addgene plasmid # 35497). The Fubi-CHR2-GFP is a plasmid that generates channel-rhodopsin2 which is sensitive to blue light and C1V1tt plasmid generates a red-shifted channel-rhodopsin that is sensitive to green light [24]. To transfect neurons, Lipofectamine LTX (catalog no. 15338500, ThermoFisher Scientific) was used with either plasmid with a typical transfection efficiency of ~10-25%. Expression for GFP or YFP-tagged transfected cultures were confirmed using fluorescence microscopy. The cultures were incubated with Lipofectamine LTX/plasmid complexes for 15 min as per manufacturer’s protocol. Post-transfection, the cultures were further incubated in media for 24-36 hours to develop appropriate expression levels prior to stimulation with OLEDs and recording with Plexon™ systems.

**Fig.3.**
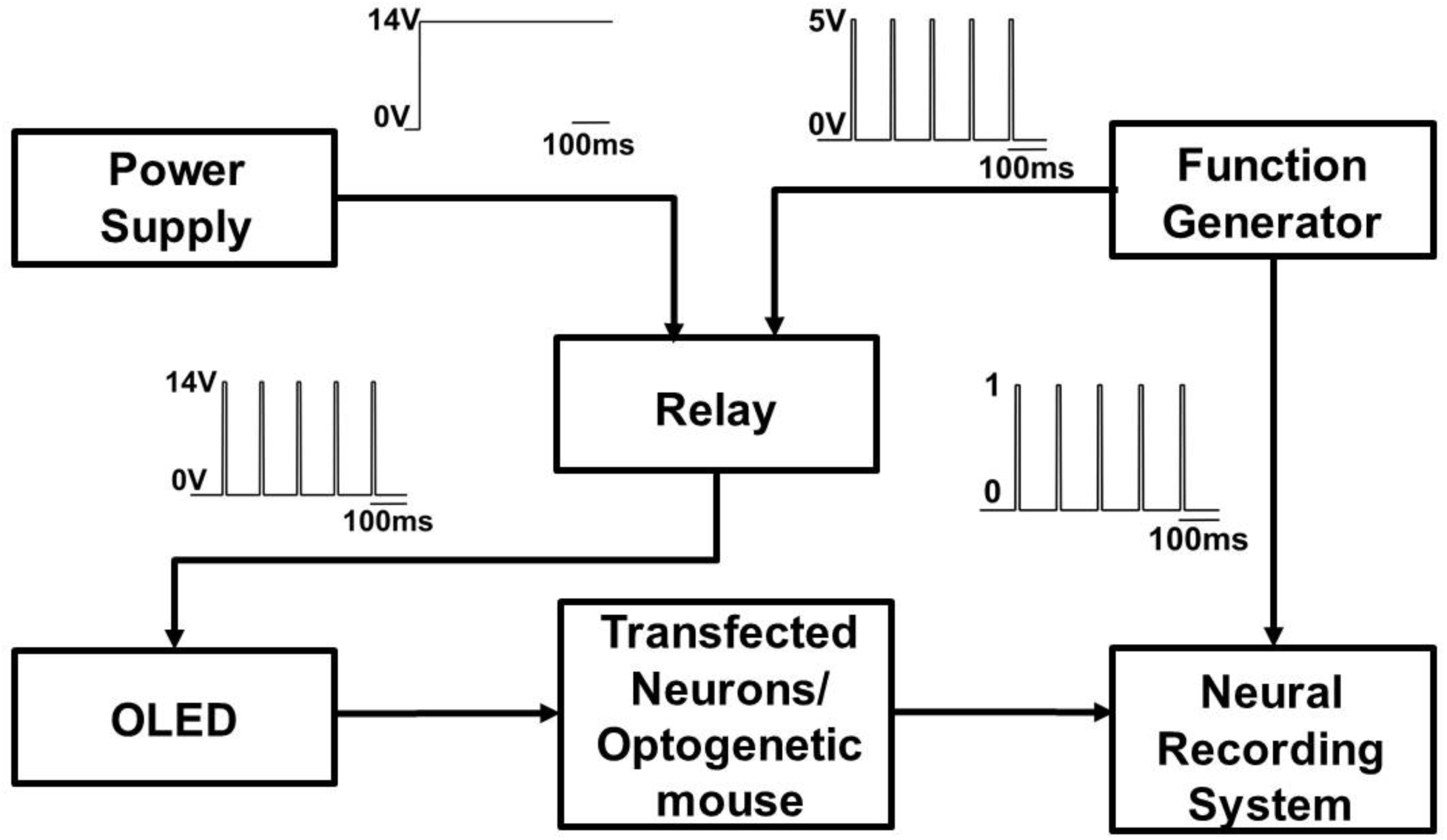
Schematic depicting optogenetic experimental setup. A DC power supply (1) generating 14 V is connected to a relay. A square wave trigger pulse from a function generator (2) with a 10% duty cycle is used to control the relay (3) to deliver a 14V pulse waveform with a 10% duty cycle to an OLED coupon (4). The pulse-driven OLED for both blue and green varieties are used to optically stimulate channelrhodopsin2 or C1V1 transfected primary mouse cortical neurons (5) cultured on an microelectrode array or an optogenetic mouse with channel-rhodopsin sensitivity. The neural activity from the stimulated neurons is recorded using a commercial multi-channel neural recording system (6). The function generator signal is fed in to the recording system and used as a stimulation event marker in neural recording analysis.

**Fig.4.**
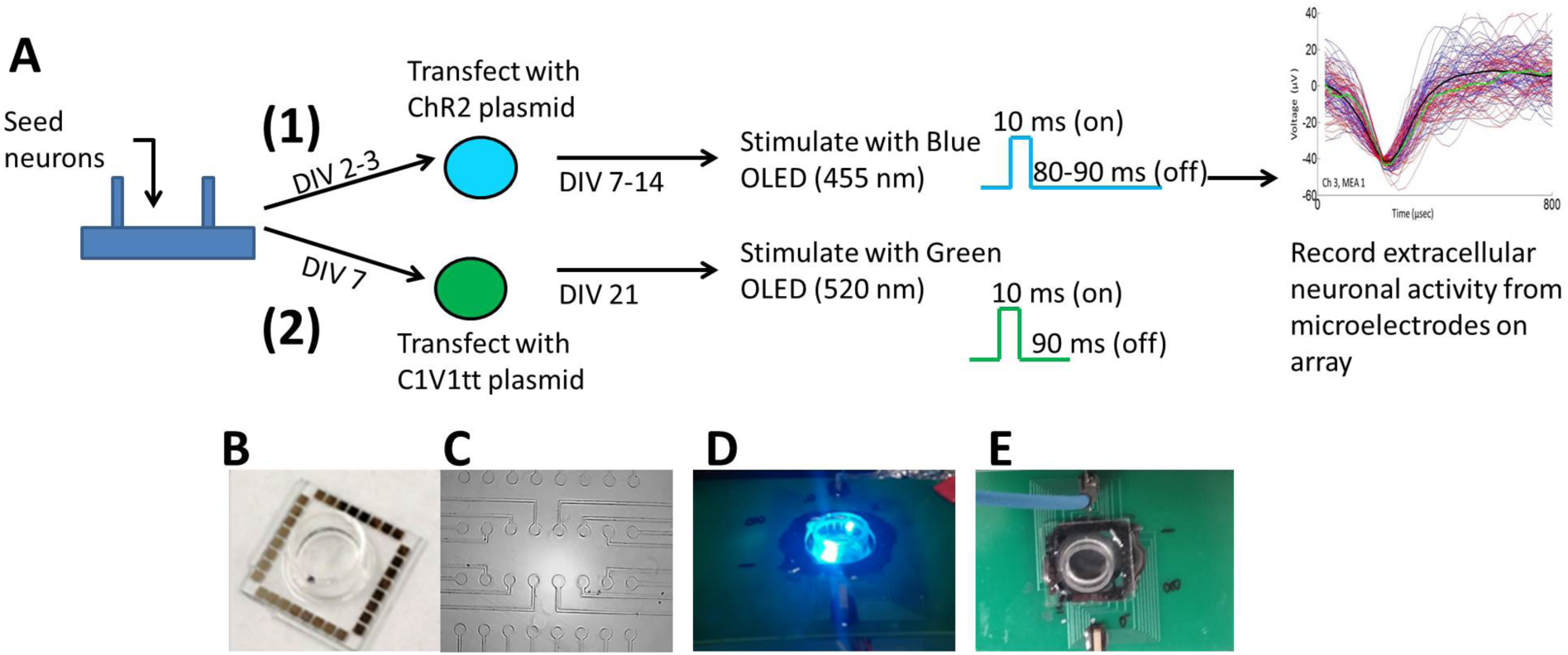
(A) In vitro experimental workflow for using pulsed OLEDs on channel-rhodopsin transfected (1) and C1V1tt transfected (2) cortical neurons on a microelectrode array (MEA). An example waveform of extracellular activity recorded from a channel on an MEA with optogenetically transfected neuron is shown. (B) Image of ITO-based MEA with (c) 32 transparent electrodes (100 µm diameter) is shown. 16 of these were capable of recording simultaneously. (D) shows an image of an example blue OLED transmitting from beneath the MEA well with (E) a recording interface connected to the MEA.

#### In vivo experimental methods

A channel-rhodopsin expressing transgenic mouse (strain: *B6.Cg-Tg (Thy1-ChR2/EYFP) 9Gfng/J*, Jackson labs) aged 12 weeks was used to test the ability of blue OLEDs to stimulate cortical neurons in an acute experiment as a proof-of-concept. All animal procedures were approved by the Institute of Animal Care and Use Committee (IACUC) of Arizona State University. The animal was induced with 42.8 mg/kg ketamine; 4.8 mg/kg xylazine; 0.6 mg/kg acepromazine administered intraperitoneally (IP) and maintained on 1-3% isoflurane during the surgical implantation of an 8-electrode -(Pt-Ir - 75 µm diameter) MEA arranged in a circle (0.5 mm radius) around an optical fiber probe (200 µm diameter) (Opto-MEA™, Microprobes, LLC., Gaithersburg, MD) and during subsequent stimulation and recording (Fig. 5). After mounting the animal on the stereotaxic frame (Kopf instruments, Tujunga, CA), the skull is exposed and craniotomies in the hindlimb motor cortex region of a mouse brain drilled was drilled ~1.5 mm lateral to midline and ~0.5 mm anterior and posterior to bregma. The Opto-MEA™ was implanted into the brain at a depth of 0.5 to 1 mm and was connected to a fiberoptic cable that collimated the OLED based light stimulation external to the mouse to the selected brain region (Fig. 5). The intensity generated by the OLED was expected to be maintained with minimal losses through the collimator and ultimate delivery through the optical fiber in the MEA.

**Fig.5.**
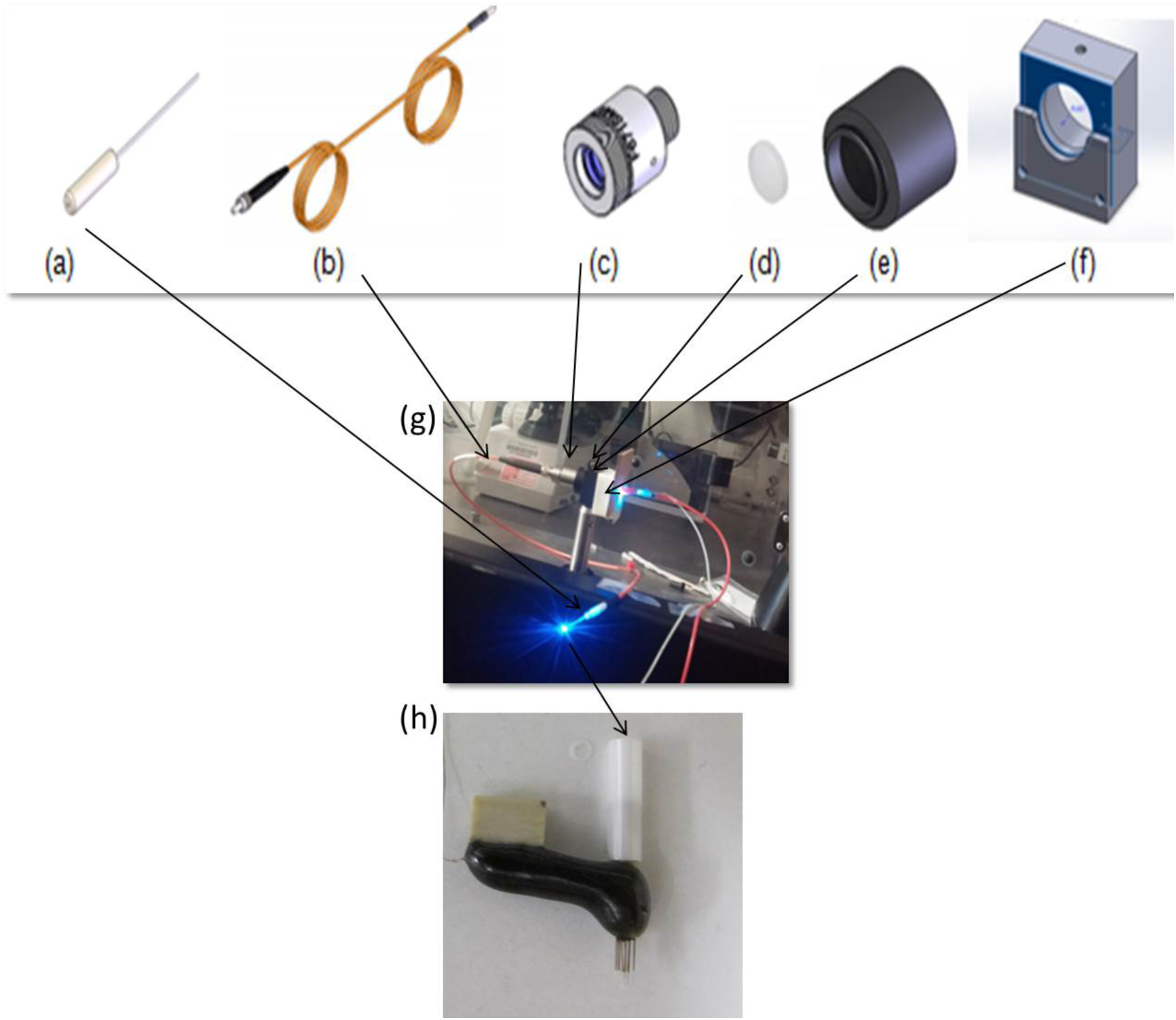
Custom designed LED light to optical fiber coupling system for in vivo photo-stimulation using pulsed OLEDs in a channel-rhodopsin expressing transgenic mouse. The following are the parts of the coupling system where (a) is the fiber optic cannula (b) fiber optic patch cable, which has a standard SMA connector that fits into the (c) collimator. The collimator subsequently connects the SMA connector and the SM05 lens tube on whose side (d) lenses are positioned inside the lens tube. (f) A 3D printed stand for the OLED flat light source is used to couple the light into the collimating system and the complete system is shown in (g). The fiber optic cable is connected to an Opto-MEA™ (Microprobes, LLC.) with 8 microelectrodes placed in a circle 500 µm in radius around a central fiber optic source. The Opto-MEA™ is implanted into the brain for simultaneous photo-stimulation and recording of neural activity.

### 2.3 Stimulation/recording and data analysis

For the in vitro MEA setup, OLEDs were placed under the transparent MEA culture dish and pulsed with 14 V, 100 msec pulse duration with 10 msec ‘on’ time and 90 msec ‘off’ time with the exception on MEA transfected with channel-rhodopsin with an ‘off’ time of 80 msec. Neural activity was recorded using Plexon™neural recording system (Plexon Inc., Dallas, TX). Noise levels were 5-8 µV during recording. Single units were sorted using Plexon™ offline software and analyzed using post-stimulus histograms (PSTHs). The PSTH plots were constructed using stimulus-evoked activity accumulated for each isolated unit on a given electrode for n=28-50 trials. Controls were taken pre-stimulus and analyzed for spontaneous, baseline activity for 5 data sets over the same time period as the post-stimulus time (100 msec) and n=28-50 trials depending on MEA. Mean ± standard deviation was calculated across the 100 msec time period for baseline activity. The mean and 95% confidence intervals for baseline activity was plotted for each MEA data set in Fig.7–10.

**Fig.6.**
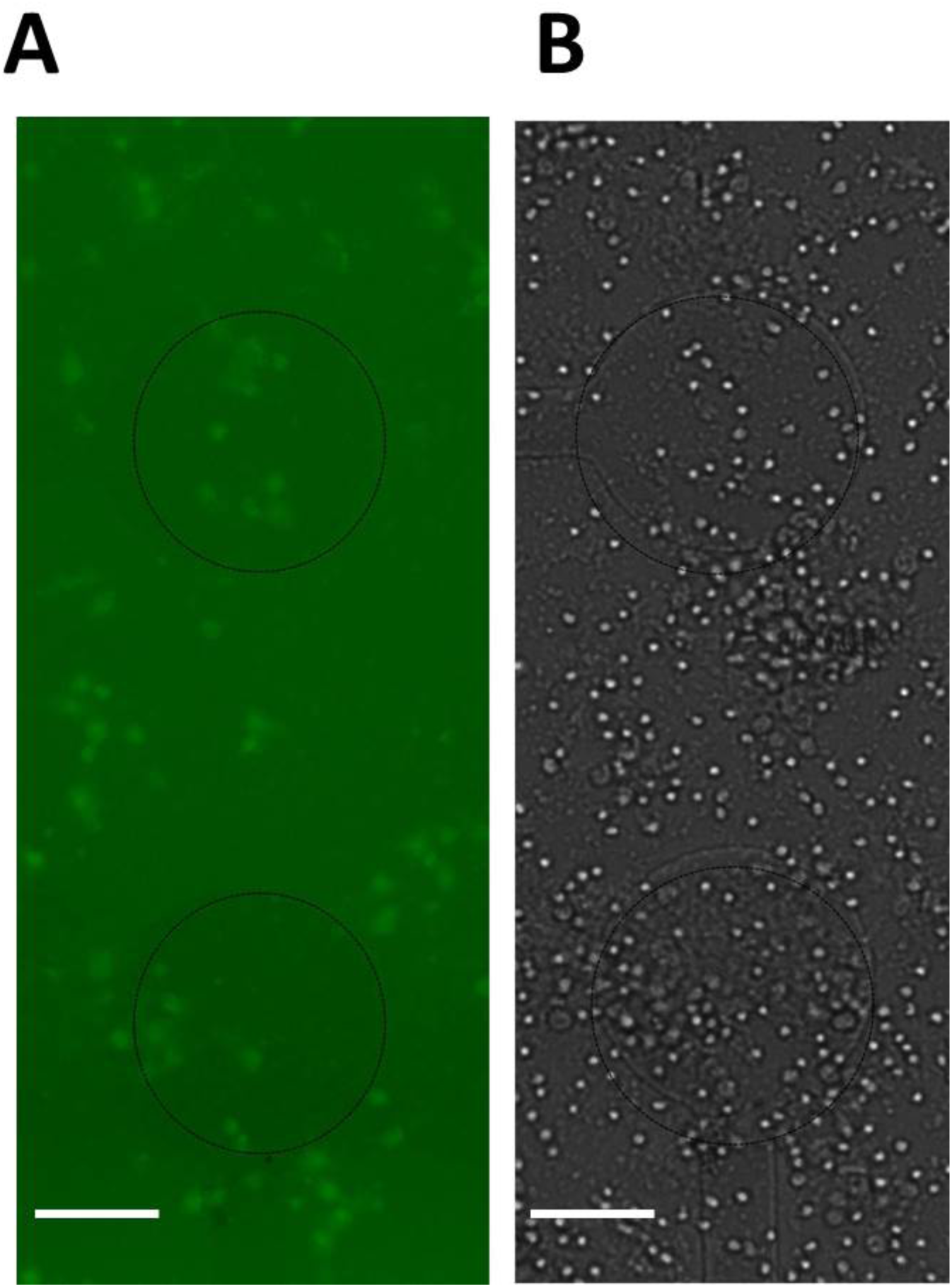
Primary cortical neurons transfected with GFP-ChR2 plasmids (A) with corresponding bright field image (B) on 2 of the microelectrodes on the MEA. Scale is 50 microns.

**Fig.7.**
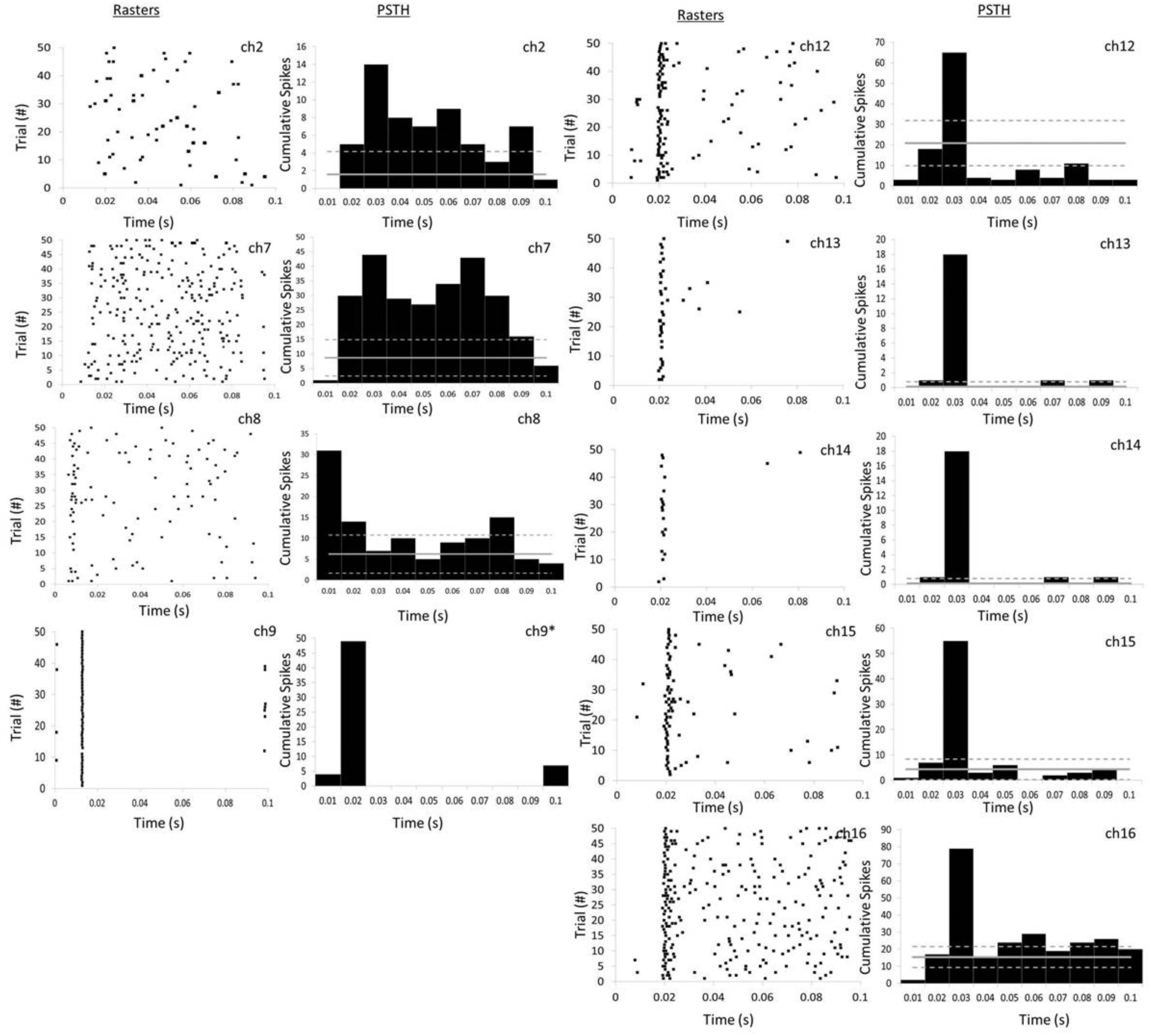
Photoactive units from an MEA with 16 recording channels transfected with channel-rhodopsin (DIV7) is shown as raster plots with corresponding post-stimulus histograms (PSTH). A total of 50 photo-stimulation trials resulted in 9 channels exhibiting 9 units responsive to OLED based photo-stimulation. All active units showed increased neural activity 10-30 msec post-stimulation. 3 of 9 units (ch 2,7,8) displayed more complex spiking behavior. The solid gray line depicts the mean baseline activity (n=5) for similar time period as photo stimulation range (0.1 s) and trial number. The dotted gray lines represent the 95% confidence intervals for baseline activity. * represents no spontaneous baseline activity, which is defined as spontaneous neural activity for a particular unit immediately prior to photo stimulation.

**Fig.8.**
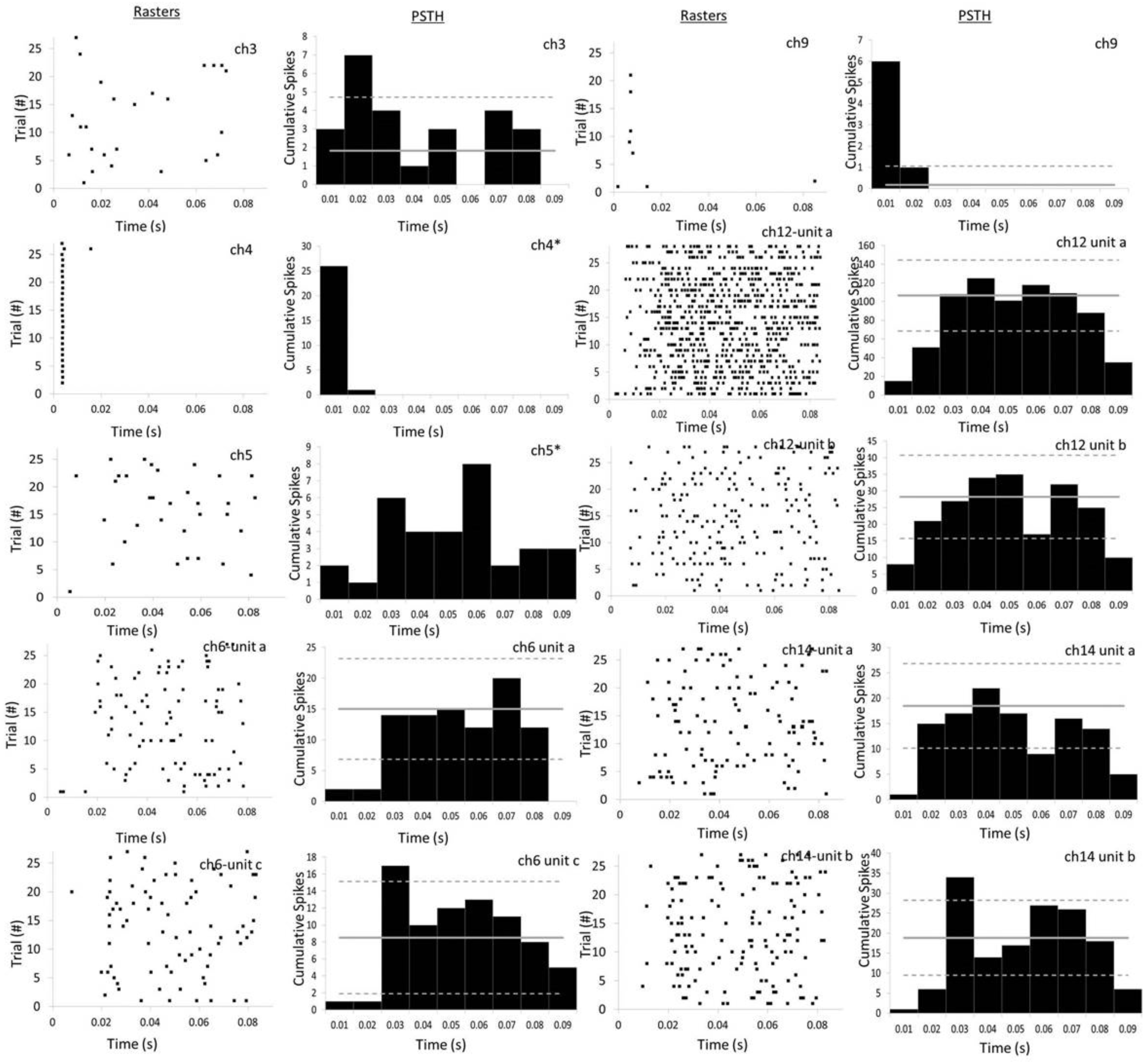
Photoactive units from an MEA with 16 recording channels transfected with channel-rhodopsin (DIV14) is shown as raster plots with corresponding post-stimulus histograms (PSTH). A total of 28 photo-stimulation trials resulted in 7 channels exhibiting 10 units responsive to OLED based photo-stimulation. Some channels had multiple units that reacted to photo-stimulation (ch6, 14). 6 of 10 active units (ch 3, 4, 5, 6c, 9, 14b) showed increased neural activity 10-30 msec post-stimulation. 6 of 10 units showed inhibitory effects (ch 6a, 6c, 12a, 12b, 14a, 14b). Interestingly, ch6 and ch14 showed 2 units displaying complex inhibitory and stimulatory activity. The solid gray line depicts the mean baseline activity (n=5) for similar time period as photo-stimulation range (0.85 s) and trial number. The dotted gray lines represent the 95% confidence intervals for baseline activity. * represents no spontaneous baseline activity, which is defined as spontaneous neural activity for a particular unit immediately prior to photo-stimulation.

**Fig.9.**
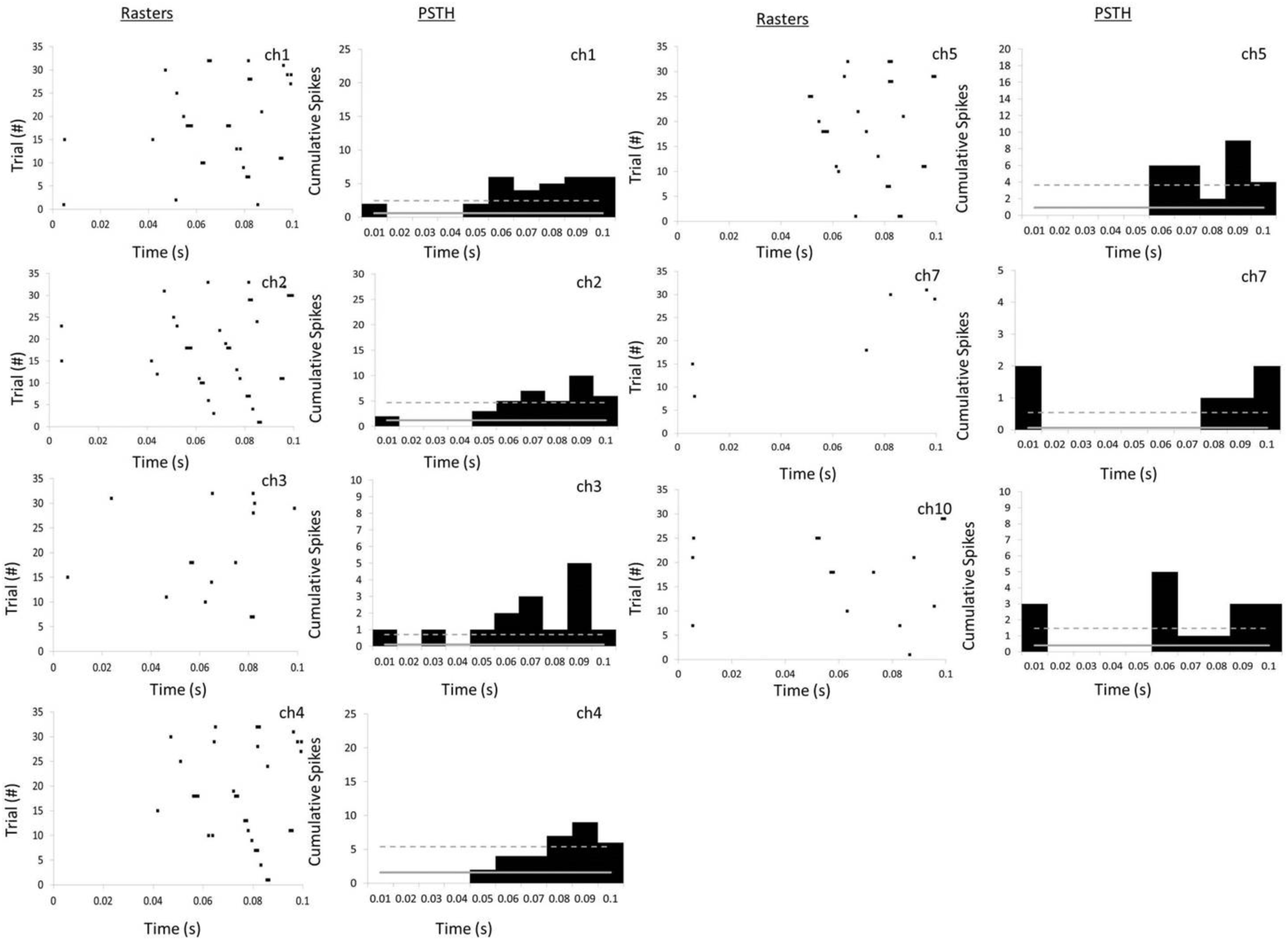
Photoactive units from an MEA with 16 recording channels transfected with C1V1tt plasmid (DIV21) is shown as raster plots with corresponding post-stimulus histograms (PSTH). A total of 32 photo-stimulation trials resulted in 7 channels exhibiting 7 units responsive to OLED based photo-stimulation. All active units showed increased neural activity ~50-60 msec post-stimulation. 2 of 7 units (ch 7, 10) displayed more complex spiking behavior with initial excitation (within 10 msec), followed by depression of activity and subsequently increase in activity 50-60 msec post-stimulation. The solid gray line depicts the mean baseline activity (n=5) for similar time period as photo-stimulation range (0.1 s) and trial number. The dotted gray lines represent the 95% confidence intervals for baseline activity, which is defined as spontaneous neural activity for a particular unit immediately prior to photo-stimulation.

**Fig.10.**
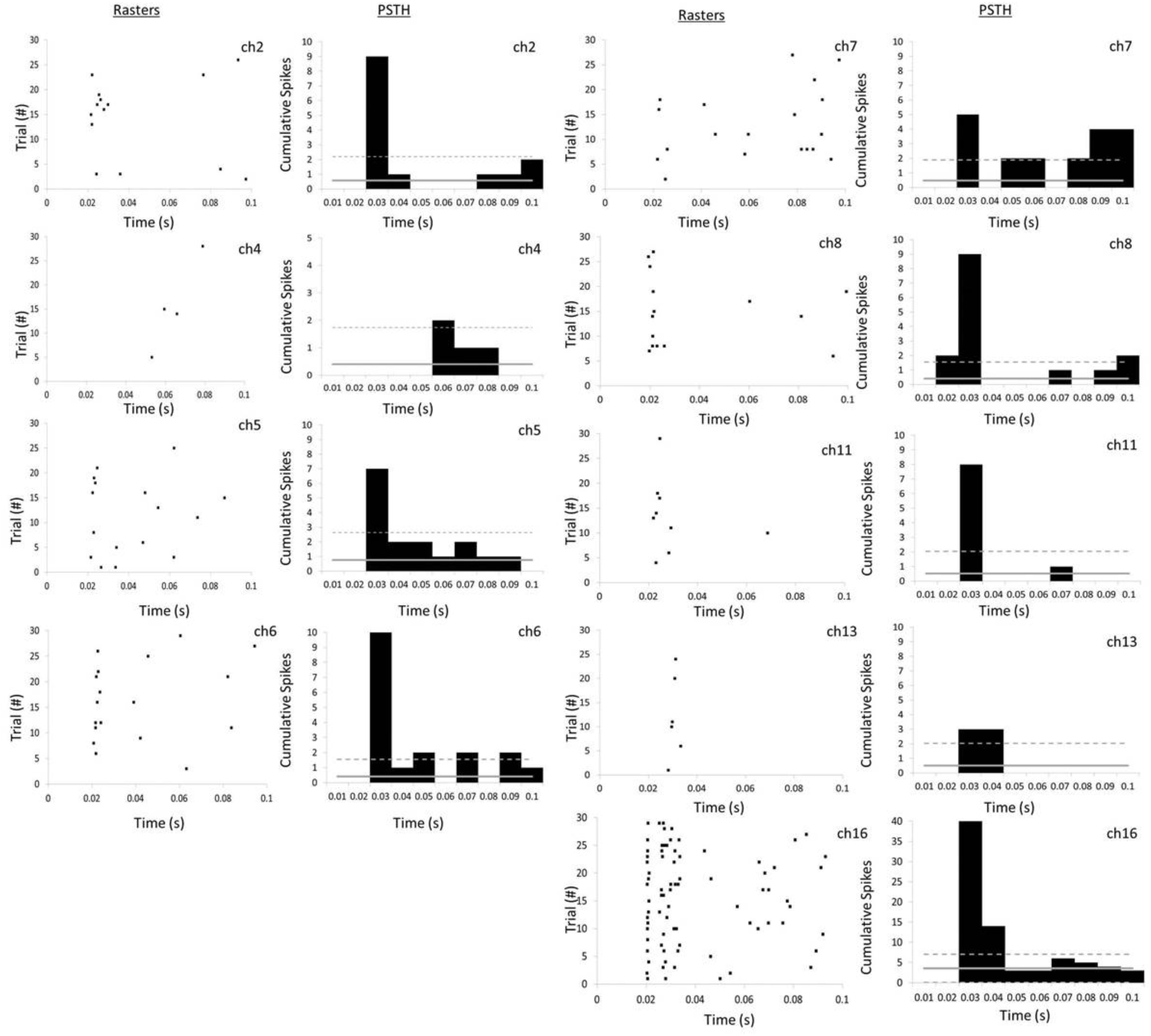
Photoactive units from an MEA with 16 recording channels transfected with c1v1tt plasmid (DIV21) is shown as raster plots with post-stimulus histogram (PSTH) analysis. A total of 30 photo-stimulation trials resulted in 9 channels exhibiting 9 units responsive to OLED based photo-stimulation. All active units showed increased neural activity ~30 msec post-stimulation. The solid gray line depicts the mean baseline activity (n=5) for similar time period as photo-stimulation range (0.1 s) and trial number. The dotted gray lines represent the 95% confidence intervals for baseline activity, which is defined as spontaneous neural activity for a particular unit immediately prior to photo-stimulation.

The in vivo OLED pulsing parameters of 14 V, 10 msec ‘on’ time and 90 msec ‘off’ time was similar to the in vitro setup. Neural recordings were obtained using TDT systems (Tucker Davis Technologies (TDT) Inc., Alachua, FL) and analyzed using custom MATLAB sorting algorithms. Noise levels were ~30 µV during recording and stimulation. Units were sorted using principal component analysis (PCA) and *k*-means statistics and analyzed for stimulus-invoked activity based on the stimulus time stamps collected using digital inputs on a separate channel. Similar to the in vitro MEA data sets, PSTHs were constructed using cumulative activity of isolated units (n=60 trials) that exhibited stimulus-evoked activity over 6 sec of data. Baseline activity for each unit was collected starting at 2 mins after termination of the stimulation protocol over a similar time period and trials, averaged over 3 baseline activity data sets, plotted as the mean and 95% confidence intervals in Fig.11.

**Fig.11.**
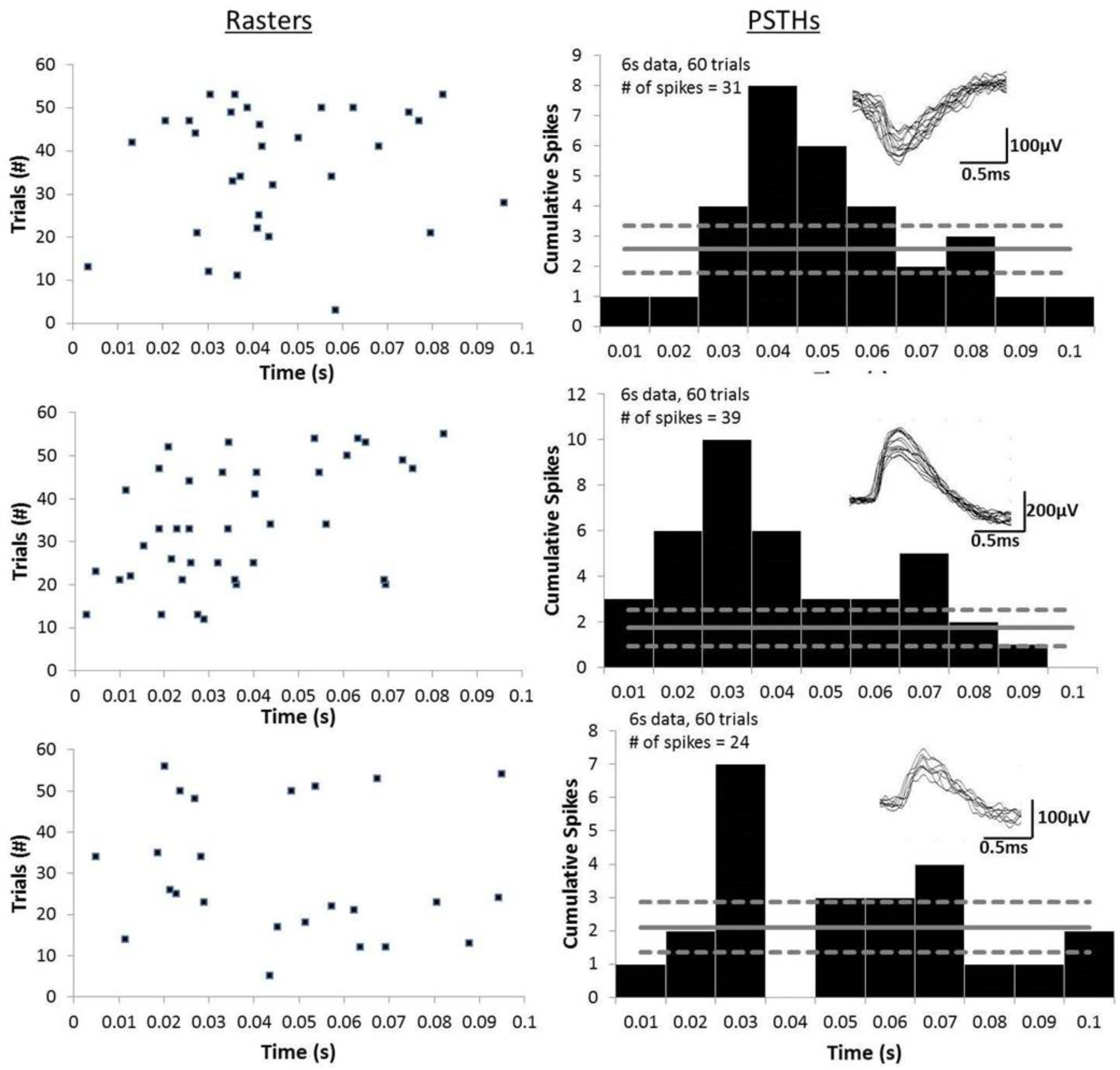
Validation of OLED based photo-stimulation in vivo. Three photoactive units from an Opto-MEA™(Microprobes, LLC.) with 8 recording channels in an optogenetic mouse (12 weeks old) expressing channel-rhodopsin is shown as raster plots with corresponding post-stimulus histograms (PSTH). A total of 60 photo-stimulation trials resulted in 3 units responsive to OLED based photo-stimulation. All active units showed increased neural activity 20-30 msec post-stimulation. The solid gray line depicts the mean baseline activity (n=3) for similar time period as photo-stimulation range (0.1s) and trial number. The dotted gray lines represent the 95% confidence intervals for baseline activity, which is defined as spontaneous neural activity for a particular unit taken at least 2 minutes post-photostimulation.

## 3. RESULTS

The primary aim of this paper was to demonstrate that OLED based light stimulators can invoke stimulus-induced activity in mammalian cortical neurons. Fig.6 shows a representative neuronal culture on a transparent MEA that has been transfected with GFP-tagged channel-rhodopsin (ChR2). The transfection efficiency was 19.6% ± 6.7% based on n=4 images. The morphology of the cells on the MEA suggested a tendency for neurons to cluster in neuronal networks, though the transfected cells were distributed uniformly across the MEA.

The raster plots and PSTH of photoactive units from ChR2 transfected MEAs are shown in Fig.7 & 8, each with 16 recording channels. In the first MEA (Fig. 7), neurons were transfected on DIV (days in vitro) 7 and neural stimulation using OLEDs and recordings were conducted on DIV 9. A total of 9 isolated neuronal units from 9 recording channels displayed OLED induced neuronal activity in over 50 photo-stimulation trials, each lasting 100 msec. As indicated in the PSTH plots, mean baseline activity taken just before the photo-stimulus application ranged from 2-20 cumulative spikes over 5 recorded snippets with a similar time period (100 msec) with same number of trials. In one channel (ch9) there was no baseline activity prior to photo-stimulation. All active units showed increased neural activity 10-30 msec post-stimulation. 3 of 9 units (ch 2,7,8) displayed more complex spiking behavior in addition to the increased level of activity poststimulation.

In the second MEA transfected with CHR2, neurons were transfected on DIV14 and neural stimulation using OLEDs (n=28 trials) and recordings were done on DIV16. A total of 10 units in 7 recording channels were isolated that were responsive to OLED stimulation. Raster plots and associated PSTHs for each unit are shown in Fig.8. Baseline activity was highly variable across channels, ranging from 2 to over 100 cumulative spikes over a similar recording time frame and number of trials. Ch 4 and 5 had no baseline activity. Overall, the isolated units on these channels displayed complex behavior in response to OLED stimulation. 4 of 10 units (labeled ch 3,4,5,9) showed increased neural activity above baseline levels at 10-30 msec post-stimulation, while another 4 of 10 units (labeled ch 6a,12a,12b,and 14a) showed inhibitory effects. Interestingly, ch 6 and ch 14 (labeled ch 6c and ch 14b) showed 2 units displaying complex inhibitory activity within the first 20 msec and subsequent excitatory activity at 30 msec post-stimulus.

OLED based photo-stimulation was also tested on 2 MEAs with neurons transfected with a red-shifted engineered channel-rhodopsin plasmid (labeled C1V1tt). Based on prior protocols [24], neurons cultured on both MEAs were transfected on DIV 21 with subsequent photo-stimulation and recordings performed on DIV 23. In the first MEA transfected with C1V1tt (Fig.9) a total of n=32 photo-stimulation trials resulted in isolation of 7 photoactive units across 7 recording channels. All active units showed increased neural activity ~50-60 msec post-stimulation. Two of 7 units (ch 7,10) displayed more complex spiking behavior, with initial stimulation (within 10 msec), followed by depression of activity and increase in activity 50-60 msec post-stimulation. In the second MEA, where neurons were transfected with C1V1tt (Fig.10), 9 channels exhibited 9 units responsive to OLED based photo-stimulation over n= 30 photo-stimulation trials. All active units showed increased neural activity ~30 msec post-stimulation. Overall, in both MEAs, the mean baseline spontaneous activity was low with <5 cumulative spikes over the same time period and number of trials across all the channels.

In the vivo rodent experiments, cumulative spikes over 60 (100 msec) trials are shown as raster plots and PSTHs for three isolated, photoactive units in Fig. 11. Typical unit amplitudes ranged from 150-500 µV. The responsive units showed significantly elevated (>95% confidence interval) neural activity 20-30 msec post-stimulation compared to baseline, spontaneous activity.

## 4. DISCUSSION

The ability of OLEDs to overcome power density limitations using pulsed mode operation and optogenetically stimulate cortical neurons in vitro and in vivo is demonstrated in this study. Prior studies have demonstrated that light intensities of 1 mW/mm^2^ from conventional LEDs are adequate to stimulate channel-rhodopsin (ChR2) transfected neurons [25][26]. As summarized in Table 1, light intensities to effectively activate ChR2 channels with 50% probability has been shown to be ~1.3 mW/mm^2^ and those for C1V1tt channels is ~0.4 mW/mm^2^ [26]. By pulsing the OLEDs at 14 V at 10 Hz, 10% duty cycle, the optical power density that can be achieved (~1 mW/mm^2^) (Fig.2) is far greater than that of the DC powered OLEDs, which cannot be operated beyond 7 V due to thermal issues. Also, the pulsed mode power density of OLEDs at 14 V appears to be sufficient to modulate the activity of neuronal networks in culture and in vivo.

**Table 1.** Characteristics of plasmids Used for OLED validation.

| <u>Plasmid</u> | <u>Typical Effect on Neuron</u> | <u>Addgene #</u> | <u>Wavelength of Peak Photocurrent</u> | <u>Effective Intensity for 50% Activation</u> |
| --- | --- | --- | --- | --- |
| ChR2 | Stimulating | 22051 | 470 nm | $1.3 \pm 0.2 \text{ mW mm}^{-2}$ |
| C1V1tt | Stimulating | 35497 | 535 nm | $0.4 \pm 0.1 \text{ mW mm}^{-2}$ |

\*Adapted from [26][24]

In comparison, recent developments using microscale GaN inorganic LEDs (iLEDs) have allowed for even higher intensities (30-100 mW/mm^2^) for microscale designs [27]. However, it is unclear how higher intensities may affect biological tissue, especially neuronal tissue which may contribute to off-target effects [28] in neuromodulation applications. Recent data would suggest that additional biocompatibility studies for use of high intensity light are warranted [29]. In addressing the need for lower intensity devices as the next generation of genetically-engineered opsins are developed, lower power requirements for stimulation are being incorporated into the opsin design (i.e. 3.0 ± 0.3 μW/mm2 for ChR2-XXL variant), allowing for more sensitivity and selectivity in activation [30]. This would be advantageous from the perspective of biocompatibility and in the future use of OLEDs as optogenetic stimulator whose design might require ever lower power consumption and enable development of novel, flexible optogenetic tools to probe neural circuits similar to those demonstrated in [21][22].

The current report utilized OLEDs that are 4 mm^2^ in area to demonstrate the ability of OLEDS to optically stimulate neuronal networks in culture and in vivo. Current iLED based technologies can achieve a microscale 50×50 µm^2^ device that potentially can be injected into tissue for wireless optogenetic experiments [27]. OLEDs demonstrated here are also scalable and are expected in future designs to achieve dimensions in the order of 25×25 µm^2^ or less using similar fabrication methodology as the reported technology. Literature also suggests the feasibility of fabricating nanoscale OLEDs [31][32]. Therefore, future application of OLED technologies will yield thinner, more flexible light sources with larger fields of view and lesser power consumption as potential advantages compared to current laser diode based light sources and competitive iLED based technologies.

An interesting result of this work is that optical stimulation of the whole culture well (all electrodes at once) induced variable, network level modulation of single units acting in concert with other neurons on MEAs. Of the MEAs transfected with ChR2 (Fig. 7,8), a total of 16 channels out of 32 channels showed photostimulus-induced behavior. Overall, 19 single units with a stimulus induced response were recorded, of which 15 units displayed an increased neural activity response as expected of ChR2 induced stimulation behavior. Three of the units in channel 9 (MEA#1, Fig. 7) and channels 4, 5 (MEA#2, Fig. 8) showed no spontaneous activity prior to photo-stimulation. Interestingly, mixed and complex spiking behavior with inhibitory and excitatory activity was seen among 9 of the units. Initial photoinduced, excitatory responses typically occurred 10-30 msec post-stimulus across both MEAs with the exception of 6 units on MEA#2 (Fig. 8, channels 6 (unit a,c), 12 (unit a,b), 14 (unit a,b)), which displayed inhibitory activity with neuronal firing rates decreasing below the lower-bound of the 95% confidence interval in the first 10-20 msec after opto-stimulation. As observed, these units also had relatively high levels of spontaneous activity (mean spikes > 8) compared to other recorded photoactive units on MEA#2. In MEA#1 (Fig.7), 33% of units (3 of 9 units on channels 2,7,8) had exhibited additional, increased neural activity 60-80 msec post-stimulus. This would suggest that an indirect effect had induced the late stimulus-driven activity from other neurons outside of the ‘listening sphere’ of the electrode but in a circuitous connection with the recording unit.

It should be noted that prior current based studies using patch-clamp and other intracellular methodologies showed that optogenetically induced changes in electrophysiology occur immediately post-stimulus. Peak currents were typically reached 3-5 msec post-stimulus for ChR2 variants [35], which is far sooner than the extracellular responses observed in this study >10-30 msec. An interesting study measuring optogenetically induced extracellular responses of DIV 7-14 cultured neurons on a MEA using a randomized spot stimuli across the culture also showed delayed post-stimulus latencies > 50 msec in select units [36]. The discrepancy between intracellular and extracellular responses could be due to variations in network excitability compared to individual neuronal excitability and the relative position of the neuron with respect to the recording electrode and point of stimulation within the network if spatial intensity variations are present within the culture well.

In MEA#2 (Fig. 8), more complex spiking behavior was seen compared to MEA#1 (Fig. 7) possibly due to the age of neuronal culture (DIV 14 for MEA#2 compared to DIV 7 for MEA#1). Also, these cortical cultures represent a mixed culture of many different neuronal types, where the network level response in a culture is highly dependent on the proportion of excitatory to inhibitory neurons (i.e GABAergic vs. glutamergic) [23][37]. Previous work suggests that 50-80% of the neurons aged > DIV 14 were sensitive to GABA, while 30-50% of the neurons were sensitive glutamate under similar culture conditions as reported in this study [23]. Typically, GABAeric neurons are characterized as fast-spiking, inhibitory interneurons, which could explain the higher levels of spontaneous activity among some units visible in MEA#2 compared to MEA#1.

The transfection efficiency for ChR2 plasmids was ~20% based on images of GFP tagged neurons (Fig.6). Representative images of the electrodes showed an average of 8-10 transfected neurons per recording electrode that potentially could respond electrophysiologically due to photo-stimulation. Assuming a random distribution of the ~3000 seeded cells/mm^2^ on the MEA and 100% viability over the number of days in culture, ~752 single neurons are expected to be recordable over a total area representative of 32 electrodes (each 100 µm in diameter). With a 20% transfection efficiency, this would hypothetically represent ~150 neurons that are potentially photoactive over 32 electrodes. However, only 19 units were recorded, where the actual number of single neurons responding to OLED stimulation represented only ~13% of the expected yield. Intensity of the light source could have played a role in the lower response since studies show that the time it takes to achieve peak current at 1 mW/mm^2^ as a measure of ChR2 channel activation would yield only 50% activation [24]. In addition, while the presence of the ChR2 gene expression in individual is identifiable with the GFP/YFP reporter gene, whether sufficient number and expression of ChR2 channels were produced for generating an action potential in a neuron is unclear. Recent studies suggest that due to low conductance, large numbers of channels need to be expressed to produce an action potential [38]. In addition, it is speculated that the levels of excitability may differ among individual neurons and their relative position or function within a neuronal circuit. In vivo studies also reflect the low number of recordable neurons, suggesting the presence of silent or dark cortical neurons [39][40]. As designs of massively parallel, multi-site high density recording based neural devices are being pursued in vivo and in vitro, the role of these neurons perhaps in terms of their relative excitability in a network needs to be better understood. The optogenetic-MEA tool might be an interesting platform to study the role of silent neurons in the future, where relative levels of excitability may be cell-addressable and programmable via optical intensity modulation.

Transfection with a red-shifted opsin encoding C1V1tt with a typical excitatory effect on a neuron, resulted in delayed activity 40-60 msec post-stimulus (Fig. 9, MEA#3) and >30msec post-stimulus (Fig. 10, MEA#4). The delayed response could be due to the slower kinetics to reach peak current compared to ChR2 variants [24]. Significantly increased activity after a delay was observed in all 16 units that exhibited photo-induced responses. Two of these units (Fig.9, MEA#3, ch 7, 10) had an initial, increased response within the first 10 msec followed by a silence and subsequent increased activity > 60 msec. These results pointed to complex, network-level modulation in these >DIV 21 cultures.

In this study, 3 units were isolated in vivo in response to photo-stimulation via a fiber optic opto-MEA system where the recording electrodes were placed ~500 µm away from the central optical fiber. The typical response occurred 20-30 msec post-stimulus (Fig. 11). Modeling studies indicate that light diffusing through tissue rapidly loses intensity to <50% of initial values within 250-500 µm around the implanted fiber-optic/light source [33][34], suggesting that only neurons closest to the fiber optic cable in the center are directly stimulated. Since the response times are slightly delayed (20-30 msec instead of immediate (<10 msec) excitation), it is speculated that the neuronal responses seen in this study are likely from indirectly stimulated neurons post-synaptic to the neurons which are directly stimulated within the network. Although, in this study, a conventional light coupling system was used to couple the OLED coupon external to the animal with the fiber optic cable (Fig. 5) implanted among cortical neurons in vivo, future iterations will involve implantable OLED thin films that will provide a direct, lens-free interface with neurons in vivo. The effectiveness of the layers of moisture barriers in protecting the OLEDs from water and oxygen (both of which can degrade OLED performance) under demanding in vivo conditions will have to be rigorously assessed in future long-term studies.

## 5. CONCLUSION

Overall, OLEDs have been shown to stimulate networks of primary cortical neurons cultured on transparent MEAs. Cortical neurons expressing ChR2 and C1V1tt opsins exhibited complex optically induced spiking behavior with excitatory and inhibitory responses in culture. The ability of OLEDs to stimulate neurons in the motor cortex was also validated in vivo using a channel-rhodopsin expressing transgenic mouse. Integration of such OLEDs with thin film transistors (TFTs) in active matrix will potentially allow the realization of implantable, flexible electronics with drastically reduced lead counts for optical stimulation of neurons in the central and peripheral nervous system.

## Acknowledgments

The authors would like to thank the Flexible Electronics and Display Center at Arizona State University (ASU), Tempe, AZ for manufacturing the prototype devices.

